# A central helical hairpin in SPD-5 enables centrosome strength and assembly

**DOI:** 10.1101/2023.05.16.540868

**Authors:** Manolo U. Rios, Bryan D. Ryder, Nicole Familiari, Łukasz A. Joachimiak, Jeffrey B. Woodruff

## Abstract

Centrosomes organize microtubules for mitotic spindle assembly and positioning. Forces mediated by these microtubules create tensile stresses on pericentriolar material (PCM), the outermost layer of centrosomes. How PCM resists these stresses is unclear at the molecular level. Here, we use cross-linking mass spectrometry (XL-MS) to map interactions underlying multimerization of SPD-5, an essential PCM scaffold component in *C. elegans*. We identified an interaction hotspot in an alpha helical hairpin motif in SPD-5 (a.a. 541-677). XL-MS data, *ab initio* structural predictions, and mass photometry suggest that this region dimerizes to form a tetrameric coiled-coil. Mutating a helical section (a.a. 610-640) or a single residue (R592) inhibited PCM assembly in embryos. This phenotype was rescued by eliminating microtubule pulling forces, revealing that PCM assembly and material strength are interrelated. We propose that interactions mediated by the helical hairpin strongly bond SPD-5 molecules to each other, thus enabling PCM to assemble fully and withstand stresses generated by microtubules.

## INTRODUCTION

Animal cells build centrosomes to organize microtubules needed for mitotic spindle assembly and chromosome segregation. Centrosomes comprise structured centrioles surrounded by a micron-scale, heterogenous protein layer termed pericentriolar material (PCM) (Conduit et al., 2015; Vasquez-Limeta and Loncarek, 2021; Woodruff et al., 2014). Self-assembly of coiled-coil-containing “scaffold” proteins, such as SPD-5 (*C. elegans*) and Centrosomin (*D. melanogaster*), form the structural foundation of PCM (Conduit et al., 2014; Woodruff et al., 2015). These proteins are recruited to the centriole surface during interphase and create a patterned layer <500 nm thick (Fu and Glover, 2012; Lawo et al., 2012; Mennella et al., 2012). In preparation for mitosis, PLK-1, Aurora A Kinase, and Spd-2/Cep192 promote growth of the PCM scaffold into a micron-scale, spherical structure (Conduit et al., 2010; Decker et al., 2011; Hannak et al., 2001; Lee and Rhee, 2011; Pelletier et al., 2004; Rathbun et al., 2020; Zhu et al., 2008). PCM becomes functional by recruiting numerous “client” proteins that nucleate microtubules and anchor them to the PCM scaffold, such as the γ-tubulin complex, TPX2, and ch-TOG (Moritz et al., 1995; Roostalu et al., 2015; Wieczorek et al., 2015; Woodruff et al., 2017). At the end of mitosis, PCM disassembles through a poorly understood mechanism. In *C. elegans*, PCM disassembly occurs in part through microtubule-mediated rupture of the PCM scaffold and PP2A-B55a Phosphatase-mediated dispersal of PCM proteins (Enos et al., 2018; Magescas et al., 2019; Mittasch et al., 2020).

PCM performs a mechanical role during mitosis by bearing forces transmitted by microtubules. Astral microtubules emanating from PCM often contact dynein motors anchored on the cell cortex; these minus-end directed motors walk toward the PCM and exert a pulling force in the range of several pN (Laan et al., 2012). Motor proteins in the mitotic spindle can also generate both pulling and pushing forces. The collective action of these microtubule-bound motors enable mitotic spindle positioning and elongation (Dumont and Mitchison, 2009). Laser ablation experiments combined with modeling indicate that the predominant forces exerted on the PCM are cortically directed, indicating that the major stress on PCM is tensile and not compressive (Farhadifar et al., 2020). Consistent with this idea, elimination of microtubules causes PCM compaction and prevents “flaring” (Enos et al., 2018; Magescas et al., 2019; Megraw et al., 2002; Rathbun et al., 2020). Although much is known about the generation of microtubule-mediated forces, very little is known about how PCM resists these forces without fracture.

The material properties of PCM are best understood in *C. elegans* embryos (Woodruff, 2021). Application of flow-based perturbations revealed that PCM transitions between a strong, ductile state to a weak, brittle state during anaphase onset (Mittasch et al., 2020). Inactivation of PLK-1 and SPD-2 (Cep192) promoted this material transition *in vivo*. These proteins reinforced SPD-5 scaffolds assembled *in vitro*, protecting them against dilution and shear stresses. Since SPD-5 is the only known PCM scaffold protein in *C. elegans*, these results suggest that PCM material properties are largely determined by tuning connections between SPD-5. Consistent with this idea, SPD-5 directly binds to microtubule-nucleating complexes; this interface likely concentrates stress, which would make SPD-5 connections critical for overall PCM strength. However, the molecular connections driving SPD-5 multimerization are poorly understood. SPD-5 (1198 a.a.) contains 9 predicted coiled-coil domains (MARCOIL, 50% threshold) and additional alpha helices separated by disordered linker regions (Alphafold, 70% confidence)(Figure 1A). An arginine-to-lysine substitution at R592 in SPD-5 strongly inhibits PCM assembly and is 100% embryonic lethal at 25°C, yet permits diminutive PCM to assemble at 16°C (Hamill et al., 2002). How this mutation affects SPD-5 structure or assembly is unknown. Pull-down and yeast-2-hybrid experiments showed an interaction between a central region containing R592 and PLK-1 phosphorylation sites (PReM-containing region; a.a. 272-734) and the CM2-like domain (a.a. 1061-1198)(Nakajo et al., 2022). While these regions are important for PCM assembly, it is not known if they contribute to PCM strength. Furthermore, since SPD-5 is sufficient to form micron-scale scaffolds, which typically require high valence (i.e., more than 3 separate binding modules)(Li et al., 2012), there are likely additional, unmapped interactions enabling SPD-5 self-assembly. Thus, it is important to comprehensively map the interactions underlying SPD-5 multimerization and resistance to microtubule-generated pulling forces.

**Figure 1.**
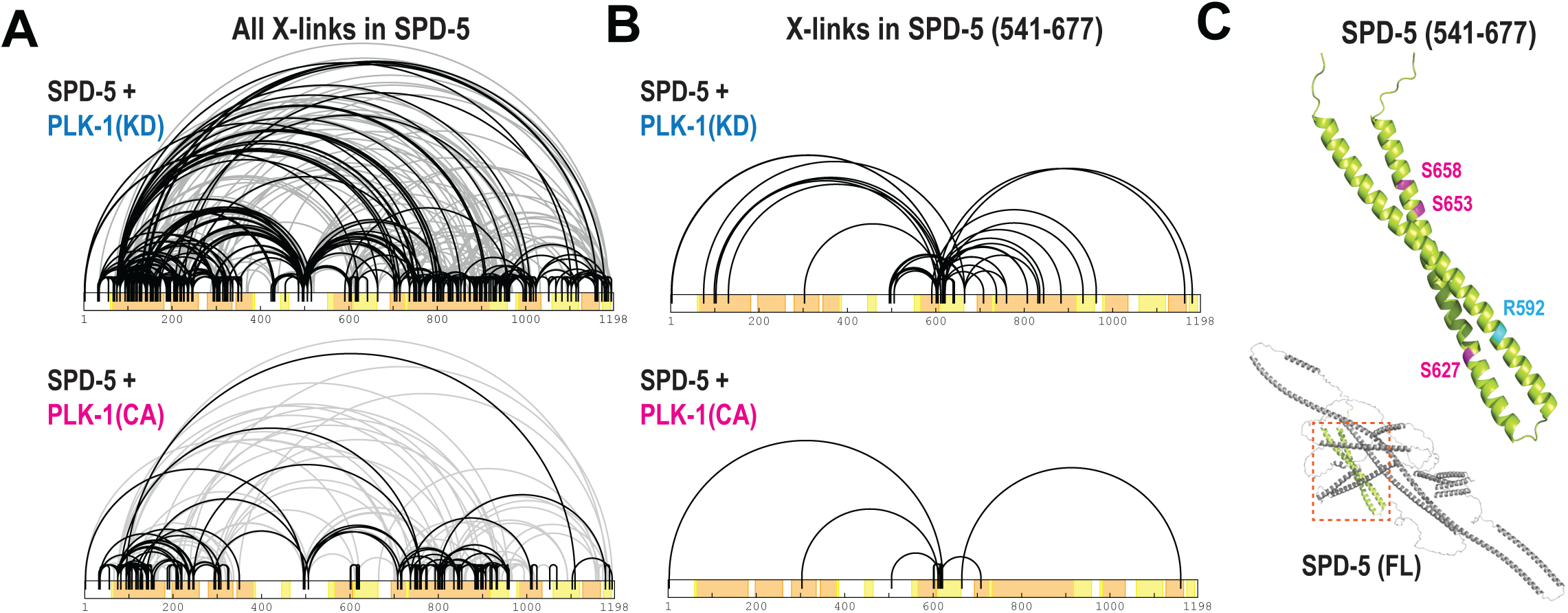
XL-MS reveals an interaction hotspot in the phospho-regulated region of SPD-5. A. Pure, full-length SPD-5 (1198 a.a.) was assembled *in vitro* in the presence of kinase dead (KD) or kinase active PLK-1 and ATP⋅MgCl_2_. The zero-length crosslinker DMTMM was added to capture contacts between lysines and glutamic or aspartic acids. Bar plots display crosslinks within SPD-5 multimers from 6 replicates; black lines represent crosslinks that appear in multiple replicates, while gray lines are crosslinks that appeared in a single replicate. SPD-5 sequence is color coded to illustrate predicted coiled-coil domains (orange), alpha helices (yellow), and disordered linkers (white). B. Crosslinks where at least one end originates in a central region of SPD-5 (a.a. 541-677). C. Alphafold predicts that the central region of SPD-5 forms an alpha helical hairpin. This motif contains Arginine 592 (blue) and PLK-1 phosphorylation sites (Serines 627, 653, and 658; magenta) that are critical for PCM assembly in *C. elegans* embryos.

Here, we used cross-linking mass spectrometry (XL-MS) to identify contact sites between SPD-5 proteins during scaffold assembly. We identified hundreds of interactions mapping to several distinct regions of SPD-5, including a PReM-localized hotspot (a.a. 541-677) containing R592 and residues required for PLK-1-mediated expansion of PCM. A companion study focuses on the interactions outside of this hotspot (Rios et al., 2023 BioRxiv). XL-MS data, combined with *ab initio* structural predictions and biophysical analyses, indicate that this region forms an alpha helical hairpin that dimerizes into a tetrameric coiled-coil with flared ends. Removing part of the second helix (Δ a.a. 610-640) or making a single substitution in the first helix (R592K) perturbed PCM assembly *in vivo*. This PCM assembly phenotype was partly rescued by eliminating microtubule-mediated pulling forces. Thus, interactions between the hairpin module are essential for generating PCM strength. These results highlight how PCM assembly and material properties are interrelated: the PCM scaffold cannot assemble unless it is strong enough to resist microtubule-dependent pulling forces. We conclude that a structured interaction between helical hairpin domains in SPD-5 undergirds the PCM scaffold.

## RESULTS

SPD-5 is essential for PCM assembly *in vivo* and sufficient to multimerize into micron-scale scaffolds capable of recruiting clients needed for microtubule aster formation i*n vitro* (Hamill et al., 2002; Woodruff et al., 2017). PLK-1 phosphorylation of SPD-5 accelerates its assembly *in vitro* and is essential for PCM expansion *in vivo* (Decker et al., 2011; Woodruff et al., 2015). To reveal how SPD-5 assembly is regulated by phosphorylation and define the molecular contacts that enable SPD-5 multimerization, we used XL-MS to map interactions found in the SPD-5 scaffold in phosphorylated and unphosphorylated states. We assembled SPD-5 into scaffolds for 2 hr in the presence of constitutively active (CA) or kinase dead (KD) PLK-1 and ATP⋅MgCl_2_. We then added the zero-length crosslinker DMTMM, which links the primary amines of lysines and the carboxylates of aspartic or glutamic acids that are <25 Å apart (cα-cα distance)(Leitner et al., 2014a). Analysis of 6 replicates per condition identified 677 (KD sample) and 289 (CA sample) crosslinks between unique residue pairs (Figure 1A). This result suggests that PLK-1 phosphorylation reduces the overall number of contacts between SPD-5 molecules. Interactions were predominantly between predicted coiled-coil motifs and alpha helical regions and not the disordered linkers. We conclude that SPD-5 assembles largely through coiled-coil-based interactions; we provide a more in-depth analysis of these interactions in our companion paper (Rios et al., 2023).

We noticed an interaction hotspot in SPD-5 (a.a. 541-677) centered around R592 and three PLK-1 phosphorylation sites that are critical for PCM expansion in *C. elegans* embryos (S627, S653, S658)(Figure 1B)(Ohta et al., 2021; Woodruff et al., 2015). We identified short-range links within this region (e.g., E602-K616, E601-K618) and long-range links with other regions of SPD-5. Phosphorylation reduced the number of contacts overall originating from this region and created a new contact with the CM2-like domain (E665-K1160)(Figure 1B). AlphaFold modeling predicts that this region contains two alpha helices connected by a loop, thus forming a helical hairpin (Figure 1C). Our data suggest that PLK-1 phosphorylation reduces non-specific, perhaps inhibitory interactions between the helical hairpin and other regions of SPD-5, thus allowing new interactions with the CM2-like domain. This interpretation is consistent with pull-down experiments showing that PLK-1 phosphorylation improves binding between purified SPD-5 fragments containing the PReM and CM2-like regions (Nakajo et al., 2022).

We next analyzed the structure and stoichiometry of the predicted hairpin region by reconstituting a fragment of SPD-5 containing residues 541-677 (Figure 2A). Circular dichroism revealed that purified SPD-5(541-677) is 72% alpha helical (Figure 2B), similar to 80% predicted by AlphaFold (confidence score >70)(Figure 1C). Mass photometry revealed that SPD-5(541-677) forms a dimer by itself and in the presence of PLK-1(KD) or PLK-1(CA) (Figure 2C,D and S1A). We verified that SPD-5(541-677) is phosphorylated by PLK-1 by gel migration shift and MS (Figure S1B). We conclude that this region is largely alpha helical and homodimerizes in a phosphorylation-independent manner.

**Figure 2.**
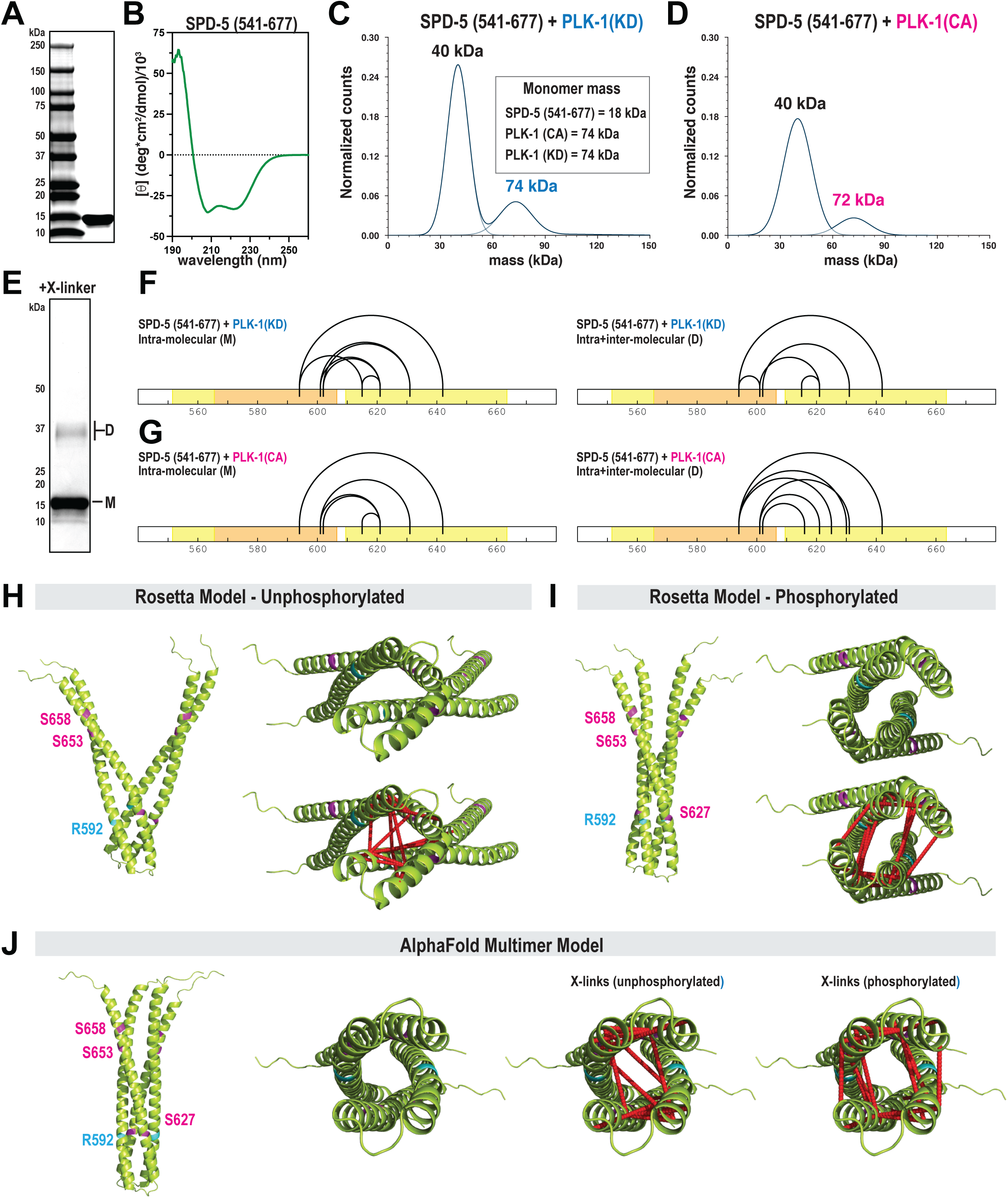
SPD-5 contains a helical hairpin motif that homo-dimerizes. A. SDS-PAGE gel showing purified SPD-5(a.a. 541-677). B. Circular dichroism of SPD-5(a.a. 541-677) reveals a strong alpha helical signature. C. 50 nM SPD-5 (541-677) was incubated with 20 nM kinase dead PLK-1 for 20 min, then analyzed by mass photometry. The inset lists the predicted monomeric masses of each protein. The calculated mass for each species is listed above the curves. D. Mass photometry of SPD-5(541-677) incubated with active PLK-1. E. Sub-saturation cross-linking of SPD-5(541-677) reveals monomeric (M) and dimeric (D) species. F. XL-MS of samples of SPD-5 (541-677) incubated with PLK-1(KD) G. XL-MS of samples of SPD-5 (541-677) incubated with PLK-1(CA). H. Best fit Rosetta model for unphosphorylated SPD-5 (541-677). Left, Top view; right, en face view. Dashed red lines indicate cross-linked residues. I. Best fit Rosetta model for phosphorylated SPD-5 (541-677). J. AlphaFold Multimer model for SPD-5 (541-677).

Having demonstrated that phosphorylation does not affect dimerization of the hairpin motif, we then tested if phosphorylation affects its three-dimensional structure. We could not build models using our XL-MS analyses of full-length SPD-5 as we could not confidently distinguish between inter- and intra-molecular contacts. We thus performed XL-MS on SPD-5(541-677) incubated with PLK-1(KD) or PLK-1(CA). By using sub-saturation amounts of crosslinker, we could generate SPD-5(541-677) species that migrate as monomers or dimers on an SDS-PAGE gel (Figure 2E), which allowed proper categorization of intra- and inter-molecular interactions in the dimer. Intramolecular interactions in SPD-5(541-677) were nearly identical in the phosphorylated and unphosphorylated states and were consistent with the hairpin configuration modeled by AlphaFold. We observed differences in crosslinking patterns between dimer samples: two links appeared only in the unphosphorylated sample (K594-E601, D615-K621), while 3 links appeared only in the phosphorylated sample (K594-E625, K594-D630, E602-K616)(Figure 2F,G). These results suggest that phosphorylation causes subtle changes in the arrangement of the two helices with respect to each other.

We then used Rosetta *ab initio* structural predictions to build a three-dimensional model of the hairpin dimer. We tested 10,000 possible docking configurations and identified best fit models based on two constraints: energy of the dimer assembly and compatibility with the experimental crosslinks, defined by the cumulative cα-cα distance of crosslink pairs for each model (Figure S1C,D). For all samples, one major model class emerged containing flared parallel dimers that form tetrameric coiled-coils near the bend in the hairpin (Figure 2H,I). For unphosphorylated SPD-5, secondary model classes emerged, which varied in positioning of the two hairpins relative to each other (Figure S1E). This result could mean that SPD-5 (541-677) can sample multiple configurations or that there are not enough crosslinks to exclude these configurations as possibilities. For phosphorylated SPD-5, all top scoring models were slight variations of the flared parallel dimer configuration (Figure 2I). Comparing the top scoring models between the unphosphorylated and phosphorylated states models revealed slight differences in dimer compaction. Thus, phosphorylation may “lock in” the tips of the hairpins and reduce flaring; however, more in-depth structural analysis is needed to test this hypothesis.

AlphaFold Multimer also predicted this region forms a tetra-helical bundle with flared ends, which appears closer to the Rosetta model for the phosphorylated state. This model was consistent with crosslinks identified in SPD-5(541-677). For all models, cross-links were <24 Å (cα-cα distance), within the acceptable range for DMTMM as demonstrated previously (Leitner et al., 2014a)(Figure S1F). Finally, the Rosetta models could account for the cross-links identified in full-length SPD-5 (Figure S1G). This result reveals the utility of using XL-MS to identify local structures within full-length SPD-5. Our results indicate that residues 541-677 in SPD-5 constitute an alpha helical hairpin that dimerizes to form a tetrameric coiled-coil with flared ends. While phosphorylation of SPD-5 does not influence dimerization of the hairpin, it could alter its connectivity and compactness.

Do the coiled-coil interactions we identified contribute to PCM strength and assembly? We addressed this question using mutational analysis and confocal microscopy of embryos. First, we used CRISPR to mutate residues 610-640, which constitute the second arm of the helical hairpin. We made the deletion in a background strain expressing SPD-5 tagged with RFP at its endogenous locus (RFP::SPD-5(WT)). Control embryos built two centrosomes that amassed spherical PCM, which cohered until rupturing and fragmenting in late anaphase, when PCM normally disassembles (Figure 3A). Embryos expressing RFP::SPD-5(Δ610-640) were viable but formed diminutive PCM that was distorted in shape (Figure 3A and S2A). RFP-positive fragments emanated from the main PCM body in *spd-5(Δ610-640)* embryos prior to anaphase, indicating premature PCM rupture. This phenotype was not due to poor expression of mutant SPD-5, as western blotting revealed similar levels of control and mutant protein (Figure S2B). In *spd-5(11610-640)* embryos, elimination of microtubule-mediated pulling forces via nocodazole treatment partially rescued the PCM assembly defect and completely rescued the PCM sphericity and premature rupture phenotypes (Figure 3A-D). These data indicate that PCM built from *spd-5(Δ610-640)* is materially weak and cannot withstand microtubule-mediated pulling forces during spindle assembly. We conclude that the integrity of the helical hairpin is critical for SPD-5 to assemble into a strong scaffold that resists microtubule-mediated forces. Given that nocodazole does not fully rescue PCM size, we conclude that the helical hairpin is also required for full PCM assembly in the absence of forces.

**Figure 3.**
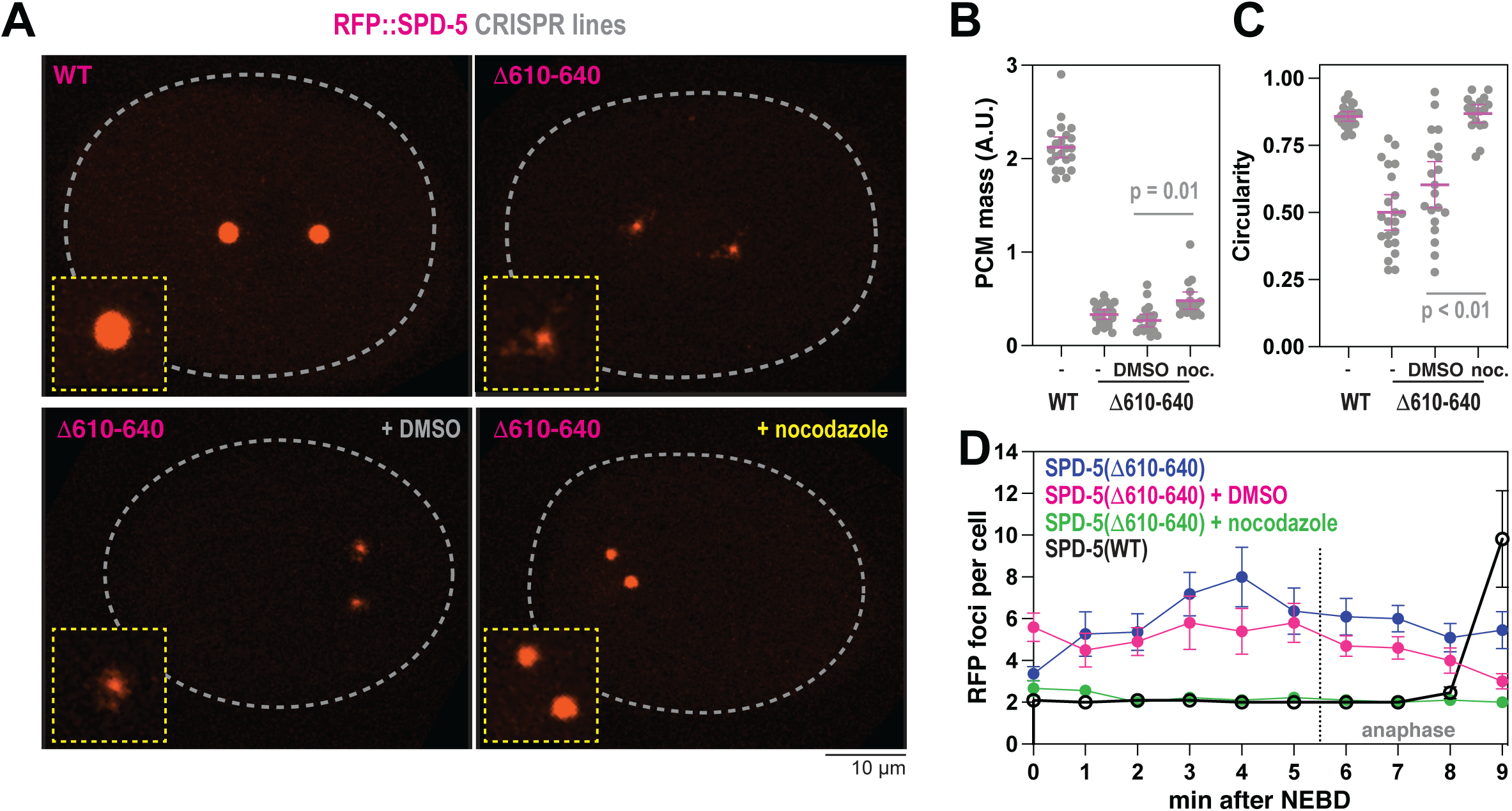
The helical hairpin motif in SPD-5 is essential for full-scale PCM assembly and strength. A. Endogenous RFP-tagged SPD-5 was modified by CRISPR to delete residues 610-640. Embryos expressing the unmodified (WT) and mutant (11610-640) RFP::SPD-5 were visualized with fluorescence confocal microscopy. Embryos were permeabilized with *perm-1(RNAi)* to allow entry of DMSO or 20 μM nocodazole. B. Quantification of fluorescence integrated density of RFP signal localized at PCM. Mean +/- 95% C.I.; N = 18-22 embryos; p value from a Kruskal-Wallis test. C. Quantification of PCM circularity. Mean +/- 95% C.I.; N = 18-22 embryos; p value from a Kruskal-Wallis test. D. PCM fragmentation from nuclear envelope breakdown onward in one-cell embryos. Mean +/- 95% C.I.; N = 9-11 embryos.

Since R592 lies within the helical hairpin of SPD-5, we hypothesized that the lethal R592K mutation disrupts PCM assembly by interfering with PCM strength. To test this hypothesis, we used immunofluorescence to visualize PCM assembly in *spd-5(or213)* embryos which express SPD-5(R592K) from the endogenous genomic locus. At 25**°**C, SPD-5 localized strongly to centrosomes in wild-type but very weakly in *spd-5(or213)* embryos, as expected (Hamill et al., 2002). Western blotting revealed similar levels of full-length SPD-5 in control and *spd-5(or213)* embryos grown at 25**°**C; thus, the PCM assembly defect is not due to lower expression levels or protein degradation (Figure S2C). Application of nocodazole rescued SPD-5(R592K) accumulation at centrosomes, although not completely to wild-type levels (Figure 4A,B). Since *spd-5(or213)* worms were generated through an EMS mutagenesis screen, it is possible that a second undetected and genetically-linked mutation could account for the PCM assembly phenotype. We thus used MosSCI to create transgenic worms expressing either wild-type or R592K versions of SPD-5 tagged with GFP (Figure 4C). These transgenes are RNAi-resistant, allowing knockdown of endogenous SPD-5 and observation of mutant phenotypes. Both control and mutant constructs expressed at similar levels (Figure S2C). In *spd-5(RNAi)* embryos, GFP::SPD-5(WT) built full-sized, functional PCM that was spherical in shape, as expected (Figure 4D,E). GFP::SPD-5(R592K) localized weakly to centrioles but was not sufficient to expand the PCM. Treatment with nocodazole significantly rescued PCM formation in GFP::SPD-5(R592K) embryos (Figure 4D,E), similar to our results with *spd-5(or213)* embryos. Thus, the R592K mutation weakens the SPD-5 scaffold, making it susceptible to premature disassembly by microtubule-mediated pulling forces. Like residues 610-640, R592 is also crucial for full-scale assembly of the PCM scaffold, independent of pulling forces.

**Figure 4.**
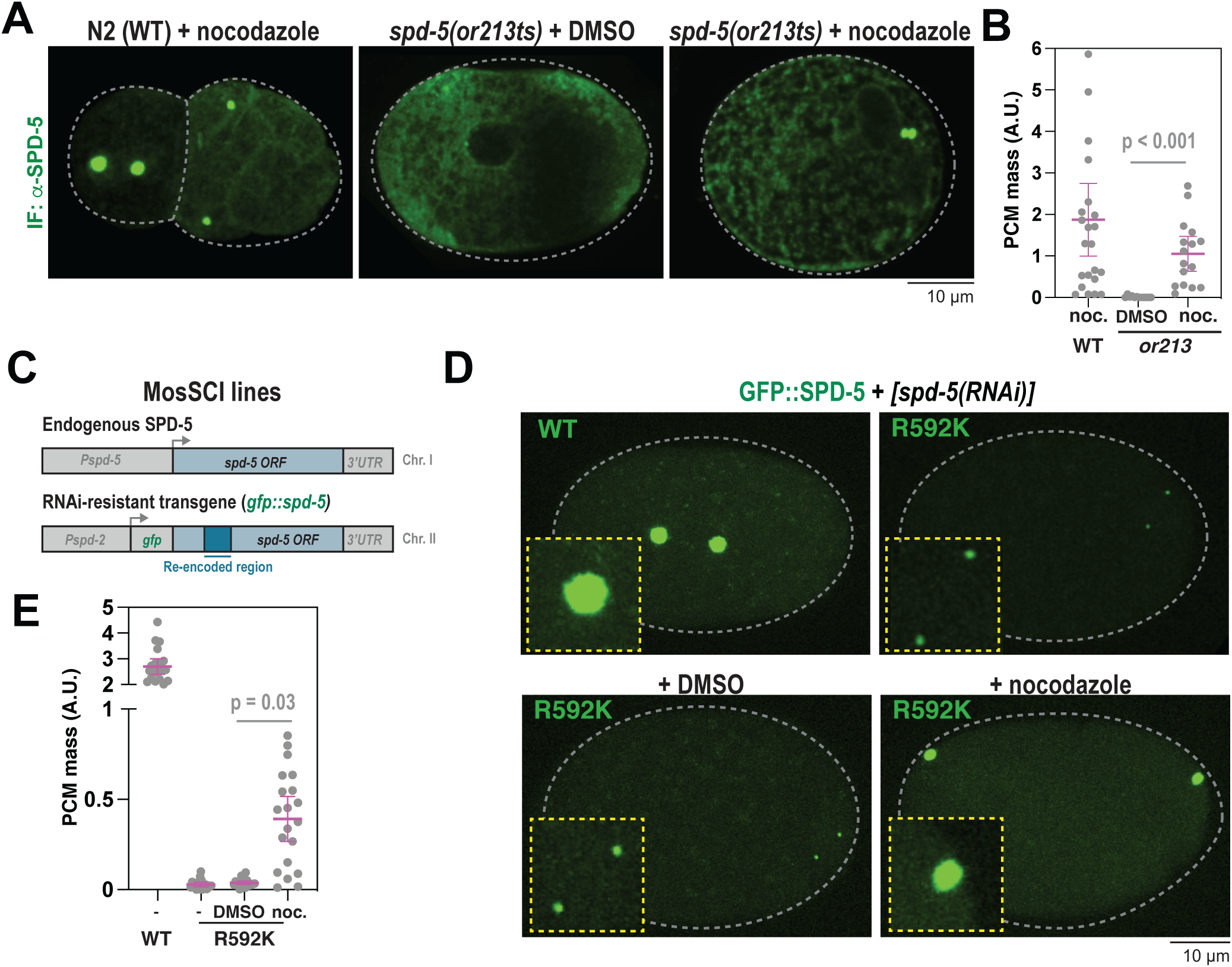
PCM defects in *spd-5(R592K)* embryos are rescued by eliminating microtubule-pulling forces. A. *spd-5(or213ts)* worms were grown at 25**°**C for 24 hr, then embryos were extracted and exposed to DMSO or 20 μM nocodazole for 2 min, then fixed. Embryos were permeabilized with *perm-1(RNAi)*. SPD-5 was detected using immunofluorescence. B. Quantification of fluorescence integrated density of SPD-5 signal localized at PCM. Mean +/- 95% C.I.; N = 12-23 embryos; p value from a Kruskal-Wallis test. C. Diagram of MosSCI-generated transgenes. Endogenous *spd-5* is expressed from chromosome I. Wild-type or mutants transgenes are expressed from chromosome II and are re-encoded to be resistant to RNAi knockdown. D. Worms were treated with RNAi against endogenous SPD-5 for 24 hr, then embryos were excised and imaged by fluorescence confocal microscopy. Embryos were permeabilized using light pressure. Insets show zoomed in images of centrosomes. E. Quantification of fluorescence integrated density of SPD-5 signal localized at PCM in (D). Mean +/- 95% C.I.; N = 18-20 embryos; p value from a Kruskal-Wallis test.

How does R592K disrupt SPD-5 scaffold assembly at the molecular level? Our structural models predict that R592 is exposed to the solvent and should only participate in hydrogen bonding with neighboring residues (Figure 5A). *In silico* analysis of energetics suggests that a change to lysine at this position should not dramatically affect stability of the hairpin dimer (Figure S3A). To test this idea, we purified SPD-5(541-677) harboring the R592K mutation and assessed its stability and dimerization capacity. Mass photometry and crosslinking revealed that both wild-type and mutant proteins are dimeric (Figure 5B and S3B). We observed dimers for both species even at the lowest concentrations of protein observable by mass photometry (10 nM), indicating a relatively high affinity. CD spectra and thermal denaturation profiles were similar between wild-type and mutant SPD-5(541-677)(Figure 5C,D). These data indicate that R592K does not detectably affect dimerization or stability of the helical hairpin, consistent with predictions from our structural models. The thermal denaturation profiles revealed a linear transition, rather than a non-cooperative transition, suggesting that the dimer lacks an extensive hydrophobic core. One interpretation is that the SPD-5(541-677) dimer has a dynamic core that is not ideally packed. This is consistent with the idea that the hairpin dimers could sample different configurations.

**Figure 5.**
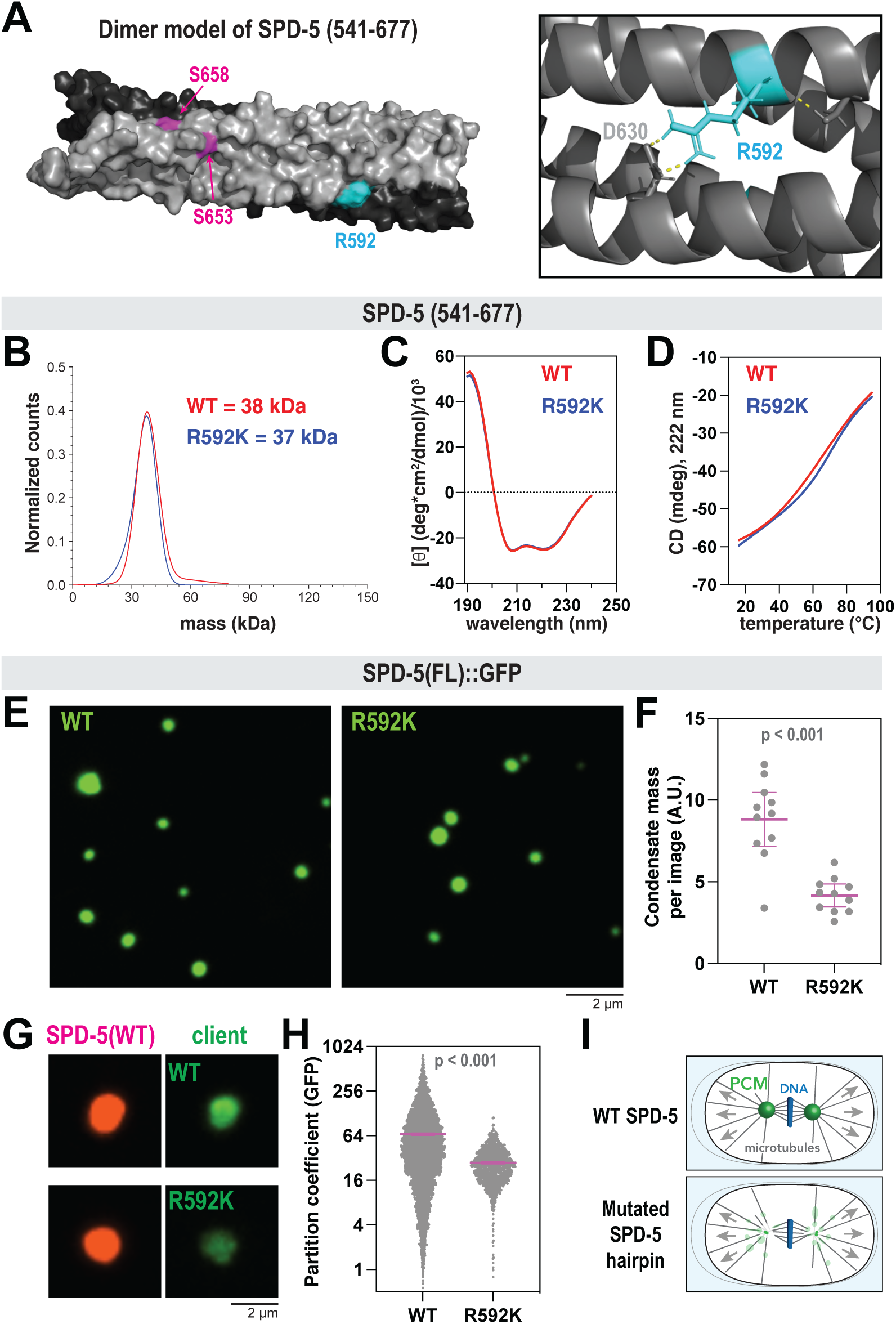
The R592K mutation disrupts SPD-5 multimerization *in vitro*. A. Surface-rendered model of a SPD-5(541-677) dimer. Key residues for one side are labeled. (Inset) Cartoon diagram of SPD-5 structure surrounding R592; dashed yellow lines represent hydrogen bonds. B. 50 nM of wild-type (WT) or mutant (R592K) SPD-5 (541-677) was analyzed by mass photometry. The calculated molecular weights are indicated. C. CD spectroscopy of WT or R592K SPD-5(a.a. 541-677). D. Thermal denaturation of WT or R592K SPD-5 (541-677) was measured by CD spectroscopy at 222 nm. E. 500 nM of purified full-length SPD-5(WT)::GFP or SPD-5(R592K)::GFP were incubated in 150 mM KCl + 7.5% PEG (MW: 3,300) for 10 min to form condensates, then imaged with fluorescence confocal microscopy. F. Quantification of (E). Mean +/- 95% C.I.;N = 11 images; p value from Mann-Whitney test. G. 1000 nM of SPD-5(WT)::RFP were assembled into condensates for 2 min, then 10 nM of client (SPD-5(WT)::GFP or SPD-5(R592K)::GFP) was added, incubated for 10 min, then imaged. H. Quantification of (G). Mean +/- 95% C.I. (N = 4702 (WT) and 1122 (R592K) condensates; p value from Welch’s t-test. I. Diagram of PCM phenotypes. Disruptions in the helical hairpin of SPD-5 causes premature PCM rupture and fragmentation, ultimately limiting PCM assembly.

Since R592 is found on the surface of the hairpin dimer, we wondered if it constitutes a binding interface to promote multimerization of full-length SPD-5. This theory is supported by our XL-MS data of full-length SPD-5, which revealed long-range interactions emanating from the helical hairpin region (Figure 1B). Therefore, we purified full-length, GFP-tagged SPD-5 (WT or R592K) and performed scaffold assembly and recruitment assays *in vitro*. 500 nM SPD-5(WT)::GFP formed micron-scale, spherical condensates in the presence of 7.5% PEG (3,000 MW), as reported previously (Woodruff et al., 2017). SPD-5(R592K)::GFP also formed spherical condensates, but the total mass of condensates per image was ∼2-fold lower compared to WT (Figure 5E,F). For the recruitment assay, we first assembled condensates using RFP-labeled SPD-5(WT) (1000 nM), then added GFP-labeled variants (10 nM). Recruitment of the R592K mutant was, on average, ∼2.5-fold lower than WT (Figure 5G,H). Thus, the helical hairpin motif is required for full-scale assembly of SPD-5 scaffolds *in vitro*. In addition, the R592K mutation directly affects assembly of the SPD-5 scaffold without affecting the local structure of the helical hairpin.

## Discussion

PCM must resist microtubule-mediated pulling forces, but how material strength emerges from the collective interactions of PCM proteins is poorly understood. Here, we identified a structural motif in SPD-5 that is essential for PCM assembly and strength in *C. elegans* embryos. This region spans amino acids 541-677 and contains residues critical for PLK-1-mediated PCM expansion. XL-MS of full-length SPD-5 combined with biophysical characterization and *in silico* modeling of a SPD-5 fragment (a.a. 541-677) suggest that this region comprises an alpha helical hairpin that dimerizes to form a flared tetrameric coiled-coil. Phosphorylation does not impede dimerization but alters the connectivity of the hairpin motif with itself and with other regions of SPD-5. Mutations in the hairpin motif caused PCM assembly defects *in vivo* that could be rescued by eliminating microtubule-mediated pulling forces. Thus, the helical hairpin acts as a “molecular linchpin” that holds the PCM scaffold together under load. The hairpin also participates in long-range homotypic interactions that assist SPD-5 multimerization. These molecular connections are crucial for PCM to achieve its full size and stay coherent when subjected to microtubule pulling forces during spindle assembly.

SPD-5 was originally identified in a screen for temperature-sensitive mutations (Hamill et al., 2002). The only identified allele (*spd-5(or213)*; R592K) prevents PCM assembly and is 100% embryonic lethal at the restrictive temperature (25°C). This mutant builds diminutive PCM at the permissive temperature (16°C). Surprisingly, we found that nocodazole rescues PCM assembly in *spd-5(or213)* embryos grown at 25°C. This result suggests that force-dependent removal of weakly attached SPD-5(R592K) ultimately prevents its accumulation around centrioles. *In vitro* assembly assays, where no force is present, revealed that SPD-5(R592K) is sufficient to self-assemble into micron-scale scaffolds but not as well as wild-type SPD-5. In a recruitment assay, where mutant protein competes with wild-type protein for docking with a pre-existing SPD-5 scaffold, wild-type SPD-5 outperformed SPD-5(R592K). Thus, we conclude that the R592K mutation primarily affects PCM scaffold strength and secondarily affects PCM scaffold assembly. Our structural models predict that R592 lies on the surface of the dimerizing hairpins. Our results support a model whereby dimerization of helical hairpins creates a binding interface that engages with other regions of SPD-5, thus reinforcing the SPD-5 scaffold. Microtubule-mediated forces diminish with decreasing temperature (Pecreaux et al., 2006), which could explain how *spd-5(or213)* embryos build small amounts of PCM and survive at 16°C.

Our results reveal how PCM assembly and material properties are interconnected: PCM must be strong enough to resist microtubule pulling forces, or else it will be ripped away from the centrosome (Figure 5I). Considering material properties could thus make sense of previous PCM assembly phenotypes in other studies. For example, expression of *pcmd-1^ts^* alleles impair SPD-5 assembly and cause visible distortions in PCM morphology, reminiscent of premature disassembly (Erpf et al., 2019). These *pcmd-1^ts^* alleles strongly diminished PCMD-1 protein levels, raising the question of how they could create a temperature-sensitive phenotype. Considering that microtubule forces decrease with temperature, these findings make sense if interpreted as a PCM weakness phenotype. We observed irregular PCM morphology and premature PCM fragmentation in embryos expressing SPD-5 lacking a.a. 610-640, which again could be rescued with nocodazole. Likewise, laser ablation of centrioles causes PCM distortion and fracture in a microtubule-dependent manner (Cabral et al., 2019). These findings suggest that departures from a spherical shape is a hallmark of PCM scaffold weakness. Thus, normal PCM flaring seen in flies (Megraw et al., 2002) and zebrafish (Rathbun et al., 2020) or PCM fragmentation phenotypes seen in human tissue culture (Yim et al., 2022 BiorXiv) could be due to weak connections between PCM scaffold proteins or between the PCM scaffold and the centriole.

Coiled-coil domains are enriched in centrosome proteins and have long been suspected to mediate assembly of the PCM scaffold (Doxsey et al., 1994; Kuhn et al., 2014; Salisbury, 2003). The functional homologs of SPD-5 in flies and humans contain 11 and 16 predicted coiled-coil domains, respectively (Centrosomin, isoform C and CDK5RAP2, isoform A). Centrosomin contains a central leucine zipper motif and CM2 domain essential for PCM assembly in fly embryos (Feng et al., 2017). In isolation, the leucine zipper forms a dimer that pairs with CM2 dimers to form a 2:2 complex of parallel coiled-coils. We see similarities with the alpha helical hairpin in SPD-5, which we demonstrate makes a tetrameric coiled-coil. While human CDK5RAP2 contains a CM2 domain (Wang et al., 2010), it is not known if this protein contains a central zipper or hairpin equivalent. Given the similarities in the general architecture of these scaffold proteins (i.e., numerous coiled-coils interspersed with linkers; CM2-like and CM1 domains), the underlying mechanism of assembly and force resistance could be the same. Testing this idea will require reconstitution and mutational analysis of PCM scaffold proteins from other species.

Finally, this study highlights the utility of *in vitro* reconstitution, XL-MS, and *ab initio* modeling to identify structural modules and reveal the mechanism of PCM scaffold assembly. Our system provides a method to comprehensively map physiologically relevant interactions that undergird PCM scaffold proteins. This current study, paired with our companion study (Rios et al., 2023), identified several uncharacterized domains in SPD-5 (the helical hairpin, a long C-terminal coiled coil, the N-terminal coiled coils) that are essential for PCM assembly and strength. We anticipate that this approach will be useful for probing the assembly of PCM scaffold proteins in other species, such as Centrosomin, PCNT, and CDK5RAP2 (Conduit et al., 2010; Jiang et al., 2021; Wang et al., 2010).

In conclusion, we identified an alpha helical hairpin motif that imparts material strength to the SPD-5 scaffold so that it can resist microtubule-mediated pulling forces. Combined with our findings in a companion study (Rios et al., 2023), our results reveal that this structured hairpin is one of many coiled-coil-based interactions that collectively enable PCM scaffold strength and assembly. These results demonstrate the importance of PCM material properties for centrosome function during embryogenesis.

## DATA AVAILABILITY

Further requests and information for resources and reagents should be directed to and will be fulfilled by the Lead Contact, Jeffrey Woodruff. Raw mass spectrometry files can be found on the MassIVE database:

Accession number MSV000090165

http://massive.ucsd.edu/ProteoSAFe/status.jsp?task=b36ed704fe8f4ed9b5100da390fe168a

Accession number MSV000090165

http://massive.ucsd.edu/ProteoSAFe/status.jsp?task=3e56f634ef304ce4b0a11366b66bbc15

Accession number: MSV000091885 https://massive.ucsd.edu/ProteoSAFe/dataset.jsp?task=fc3b993f37c34c89b5028d93bdd58ab5

## Supporting information

Supplemental Data Set 1

Supplemental Data Set 2

Supplemental Data Set 3

Supplemental Data Set 4

## ACKNOWLEDGEMENTS

We would like to thank Bruce Bowerman for providing strains; Craig Mello for providing reagents for transgenic worm construction; the Macromolecular Biophysics Resource Facility at UT Southwestern for help with CD spectroscopy and mass photometry; the Proteomics Core Facility at UT Southwestern for mass spectrometry; Jesse Bucksot and Sofia Bali for help with developing MATLAB analysis scripts.

## AUTHOR CONTRIBUTIONS

M.U.R. performed and analyzed *in vivo* experiments involving *C. elegans* embryos, purified proteins, and performed and analyzed XL-MS experiments. B.D.R. and L.A.J. analyzed cross-linking data. N.F. performed baculovirus-mediated protein expression and immunofluorescence. J.B.W. performed *in vitro* assays, optimized XL-MS protocols, and analyzed data. L.A.J. performed Rosetta simulations. J.B.W. wrote the manuscript.

## MATERIALS AND METHODS

### Experimental model and subject details

For expression of recombinant proteins (listed in Table S1) we used ArcticExpress (DE3) bacteria (Agilent) or SF9-ESF *S. frugiperda* insect cells grown at 27**°**C in ESF 921 Insect Cell Culture Medium (Expression Systems) supplemented with Fetal Bovine Serum (2% final concentration). *C. elegans* worm strains were grown on nematode growth media (NGM) plates at 16-25**°**C, following standard protocols (www.wormbook.org). Worm strains used in this study are listed in Table S2 and created using MosSCI (Frokjaer-Jensen et al., 2008) or CRISPR (Paix et al., 2015).

### Prediction of coiled-coil motifs and secondary structure

Coiled-coil motifs in SPD-5 were predicted using MARCOIL at 50% threshold, an MTK matrix, and high transition probability (https://toolkit.tuebingen.mpg.de/tools/marcoil). Secondary structure was predicted using Alphafold (Jumper et al., 2021; Varadi et al., 2022).

### Ab initio modeling

Initial conformation of the hairpin conformation of the SPD-5 541-677 fragment was generated using AlphaFold from full-length SPD-5 (Jumper et al., 2021; Varadi et al., 2022). The dimerization of this fragment was evaluated using two parallel strategies: AF multimer module (https://doi.org/10.1101/2021.10.04.463034) and Rosetta docking (Chaudhury et al., 2011; Gray et al., 2003). For Rosetta modeling of the dimer, 10,000 models were generated in the absence of restraints using the low resolution centroid mode followed by high resolution refinement (Chaudhury et al., 2011). Additionally, we compared the energetics of the WT and R592K SPD-5 fragments (541-677) to evaluate the effect of the mutation on the stability of the dimer ensemble. The ensemble of dimeric models was evaluated against crosslinks obtained from the full-length and fragment SPD-5 XL-MS datasets by computing the sum of the distances for all experimentally observed cα-cα distance XL-MS pairs. Because no symmetry was imposed on the dimer generation the sum of distances were computed for both geometries to test the symmetry of the interfaces. Final models were compared that exhibited low computed assembly energies and low cumulative distance between crosslink pairs.

### Protein purification

All expression plasmids are listed in Table S1. Full-length SPD-5 and PLK-1 constructs were expressed and purified as previously described (Woodruff and Hyman, 2015; Woodruff et al., 2015). For newly made proteins, the full-length *spd-5* coding sequence was inserted into a baculoviral expression plasmid (pOCC27, pOCC28, or pOCC29) using standard restriction cloning. Baculoviruses were generated using the FlexiBAC system (Lemaitre et al., 2019) in SF9 cells. Protein was harvested 72 hr post infection during the P3 production phase. Cells were collected, washed, and resuspended in harvest buffer (25 mM HEPES, pH 7.4, 150 mM NaCl). All subsequent steps were performed at 4**°**C. Cell pellets were resuspended in Buffer A (25 mM HEPES, pH 7.4, 30mM imidazole, 500 mM KCl, 0.5 mM DTT, 1% glycerol, 0.1% CHAPS) + protease inhibitors, then lysed using a dounce homogenizer and sonication (20% amplitude, 5 sec ON, 15 sec OFF, for 90 sec). Proteins were bound to Ni-NTA (Qiagen), washed with 10 column volumes of Buffer A, eluted with 250 mM imidazole. The eluate was then bound to amylose resin (NEB), washed with 5 column volumes of Buffer C (25 mM HEPES, pH 7.4, 500 mM NaCl, 0.5 mM DTT, 1% glycerol, 0.1% CHAPS). Protein was eluted by adding PreScission protease, incubating overnight, and then passed over Glutathione Sepharose 4B (Sigma) to remove the Precission protease. Eluted protein was then concentrated using 3K-50K MWCO Amicon concentrators (Millipore). All proteins were aliquoted in PCR tubes, flash-frozen in liquid nitrogen, and stored at −80**°**C. Protein concentration was determined by measuring absorbance at 280 nm using a NanoDrop ND-1000 spectrophotometer (Thermo Scientific).

For SPD-5 (541-677), a codon-optimized gene block was cloned into the 6xHis_1B LIC plasmid (gift from Scott Gradia, UC Berkeley) and transformed into Arctic Express bacteria (Agilent). Expression was induced with 0.3 mM IPTG for 16 hr at 20**°**C. Cell pellets were resuspended in Buffer A + protease inhibitors, then lysed using an Emulsiflex followed by sonication (40% amplitude, 5 sec ON, 15 sec OFF, for 90 sec). Lysate was centrifuged at 24,000 rpm in a JA-25 rotor for 30 min. Supernatant was passed over Ni-NTA agarose, washed, and eluted with 250 mM imidazole. Eluate was concentrated with an 10K MWCO Amicon concentrator (Millipore), filtered, then passed over a Superdex 75 Increase size exclusion column (Cytiva). Fractions containing SPD-5(541-677) were concentrated and aliquoted in PCR tubes, flash-frozen in liquid nitrogen, and stored at −80**°**C. Protein concentration was determined by measuring absorbance at 205 nm using a NanoDrop ND-1000 spectrophotometer (Thermo Scientific).

### Cross-linking mass spectrometry of SPD-5

A cumulative list of cross-linked pairs can be found in Supplemental Data Set 1. To achieve reliable results, we found it was necessary to start with completely dephosphorylated SPD-5. Purification of dephosphorylated SPD-5 was achieved through modification of our standard protocol. In short, insect cell lysates were passed through 4mL of Ni-NTA beads twice, washed five times with buffer 1 (25mM HEPES, 500mM NaCl, 30mM imidazole, 1% glycerol, 0.1% CHAPS, pH 7.4), then twice with buffer 2 (150mM KCl, 25mM HEPES, pH 7.4) at 4°C. Ni-NTA-bound SPD-5 was incubated for 1 hr at room temperature in dephosphorylation buffer (1X PMP buffer (NEB) + 1mM MnCl_2_, + 40,000 U of lambda phosphatase (400,000 units/mL, NEB). Beads were then washed twice with buffer 1 at 4°C. Dephosphorylated SPD-5 was eluted from the Ni-NTA beads using 15mL of Buffer 3 (25mM HEPES, 500mM NaCl, 250mM imidazole, 1% glycerol, 0.1% CHAPS, pH7.4). SPD-5 was then bound to 500 μL MBP-trap beads (Chromotek) and the column was washed 3X with buffer 4 (25mM HEPES, 500mM NaCl, 1% glycerol, 0.1% CHAPS, pH7.4). Dephosphorylated SPD-5 was eluted from the MBP-trap beads by overnight incubation in buffer 4 + 100μL of PreScission protease (Acro Biosystems; 1mg/mL) at 4°C. Eluted protein was further purified and stored as described above. Dephosphorylation of SPD-5 was confirmed by PTM identification using mass spectrometry showing effective removal of 99.8-100% of phosphates.

Crosslinking reactions were prepared at room temperature with 1μM dephosphorylated SPD-5, 1μM PLK-1 (KD/CA), 0.2 mM ATP, 10 mM MgCl_2_, 150mM KCl, 25mM HEPES, pH7.4 and 0.5mM DTT. Samples were incubated for 2 hr at room temperature followed by addition of 8 mM DMTMM for 45min at room temperature (shaking at 300 rpm). To quench the reaction, we added 50 mM ammonium bicarbonate for 15min at room temperature (shaking at 300rpm). Samples without cross-linker were used for mass spectrometry PTM analysis to identify phosphorylated sites (Supplemental Data Set 2).

Samples were run on an SDS-PAGE gel to separate cross-linked species. Bands corresponding to monomeric or multimeric protein were excised from the gel, then digested overnight with trypsin (Pierce), reduced with DTT, and alkylated with iodoacetamide (Sigma). Samples were cleaned using solid-phase extraction with an Oasis HLB plate (Waters), then injected into an Orbitrap Fusion Lumos mass spectrometer coupled to an Ultimate 3000 RSLC-Nano liquid chromatography system. Peptides were separated using a 75 μm i.d., 75-cm long EasySpray column (Thermo) and eluted with a gradient at a flow rate of 250 nL/min from 0-5% buffer B over 1 min, 5%-40% B over 60 minutes, 40%-99% over 25 minutes, and held at 99% B for 5 minutes before returning to 0% B for column equilibration. Buffer A contained 2% (v/v) ACN and 0.1% formic acid in water, and buffer B contained 80% (v/v) ACN, 10% (v/v) trifluoroethanol, and 0.1% formic acid in water. The mass spectrometer operated in positive ion mode with a source voltage of 1.5-2.4 kV and an ion transfer tube temperature of 275°C. MS scans were acquired at 120,000 resolution in the Orbitrap and up to 10 MS/MS spectra were obtained in the ion trap for each full spectrum acquired using collision-induced dissociation (CID) for ions with charges 3-7. Dynamic exclusion was set for 25 s after an ion was selected for fragmentation.

For data analysis, each Thermo.raw file was converted to .mzXML format for analysis using an in-house installation of xQuest (Leitner et al., 2014b). Score thresholds were set through xProphet(Leitner et al., 2014b), which uses a target/decoy model. The search parameters were set as follows. For zero-length cross-link search with DMTMM: maximum number of missed cleavages = 2, peptide length = 5–50 residues, fixed modifications carbamidomethyl-Cys (mass shift = 57.02146 Da), mass shift of cross-linker = −18.010595 Da, no monolink mass specified, MS^1^ tolerance = 15 ppm, and MS^2^ tolerance = 0.2 Da for common ions and 0.3 Da for cross-linked ions; search in enumeration mode. For grouping heavy and light scans (DSS cross-links only): precursor mass difference for isotope-labeled succinimides = 12.07573 Da for DSS-h_12_/d_12_; maximum retention time difference for light/heavy pairs = 2.5 min. Maximum number of missed cleavages (excluding the cross-linking site) = 2, peptide length = 5–50 aa, fixed modifications = carbamidomethyl-Cys (mass shift = 57.021460 Da), mass shift of the light crosslinker = 138.068080 Da, mass shift of mono-links = 156.078644 and 155.096428 Da, MS^1^ tolerance = 10 ppm, MS^2^ tolerance = 0.2 Da for common ions and 0.3 Da for cross-link ions, search in enumeration mode. Samples were also re-analyzed accounting for mass shifts due to the presence of phospho-serines and phospho-threonines. The false discovery rates (FDR) for all experiments were <20% (full length SPD-5) and <5% (541-677) at the link level (see Supplemental Data Sets 3 and 4 for further information about cross-linked peptide pairs). For XL-MS of SPD-5(541-677), only the top 10% of cross-links (based on protein ID score) were used for modeling and display in Figure 2, making the final FDR <1%.

### Mass Photometry

500 nM SPD-5(541-677) was centrifuged to remove potential aggregates, then diluted into PBS to a final concentration of 50 nM on a clean glass coverslip. Protein was analyzed using a TwoMP mass photometer (Refeyn), using four species BSA (monomer, dimer, trimer, tetramer) to create a molecular weight standard curve.

### RNAi treatment

RNAi was done by feeding. The *spd-5* feeding clone targets a region that is re-encoded in our MosSCI transgenes (Woodruff et al., 2015). Bacteria were seeded onto nematode growth media (NGM) supplemented with 1 mM isopropyl β-D-1-thiogalactopyranoside (IPTG) and 100 µg mL^-1^ ampicillin. L4 hermaphrodites were grown on feeding plates at 23°C for 24 hours. For *perm-1(RNAi),* worms were grow on NGM + 0.1 mM IPTG at for 23°C 16 hours (Carvalho et al., 2011).

### Western Blotting

60 adult worms were picked and transferred to blank plates for 20 min to remove bacteria off their bodies. Worms were then moved to PCR tubes containing 10µL of mili-Q water to which 10µL of SDS loading buffer was added. Each sample was placed in a pre-warmed heat-block (95°C) for 5 minutes. 5µL per sample was then loaded into three 4-20% mini-protean TGX gels and ran for 35min at 200V. Proteins from each gel was transferred to a nitrocellulose membrane using a Trans-blot turbo transfer for high molecular weight proteins (10 minutes). Membranes were incubated in blocking buffer consisting of 1X TBS-T + 3% Blotting-Grade Blocker (BioRad) shaking at room temperature for 1 hr. Membranes were then washed three times with fresh 1X TBS-T and incubated shaking with primary antibodies over night at 4°C. Primary antibody was washed three times with fresh 1X TBS-T and incubated shaking with secondary antibodies at room temperature for one hour. Each membrane was then incubated in ECL reagent (Thermo Scientific SuperSignal West Femto) for 5 minutes and imaged with a ChemiDoc Touch Imaging System. <u>Primary Antibodies:</u> Rabbit anti-SPD-5 (1:1,000, clone 758, Dresden Antibody Facility). <u>Secondary Antibodies (1:50,000):</u> HRP conjugated Goat anti-Rabbit IgG (1mg/mL) (Invitrogen, # 65-6120, Lot # J276300).

### Embryo viability assay

10 L4 worms picked and transferred to OP50 plates. 24 hours later, individual adult worms were transferred to individual plates (10 worms per strain) and allowed to lay eggs for 6 hr. Adult worms were then removed from the mating plates and eggs were manually counted. Hatched worms were then counted 24 hr later. Plate viability is reported as number of worms hatched divided by number of eggs laid. Strain viability is reported as the average viability of 10 plates.

### Generation of *C. elegans* embryos expressing *spd-5* transgenes

*C. elegans* worm strains used in this study were created using MosSCI (Frokjaer-Jensen et al., 2008) and based on constructs made previously (Woodruff et al., 2015).The GFP::SPD-5(R592K) mutant was created by modifying an existing MosII compatible plasmid (pOD1021_*pSpd-2::gfp::spd-5(RNAi-resistant)::spd-5_3’UTR*). Mutated sequence was synthesized by PCR and used to replace a section of the wild-type sequence. Plasmids were purified using a NucleoBond Xtra Midi Prep Kit (Macherey Nagel), combined with co-injection plasmids, and injected into strain EG6699 (tt5605, Chr II). After one week, worms were heat-shocked for 3 hr at 35℃ to kill worms maintaining extrachromosomal arrays. Moving worms without fluorescent co-injection markers were selected as candidates. Sequencing was used to confirm transgene integration.

### Generation of CRISPR-modified *C. elegans* mutants

*C. elegans* worms expressing tagRFP::SPD-5 at the endogenous spd-5 locus (gift of J. Feldman) were modified by CRISPR-Cas9 to delete the 93 bp sequence encoding a.a. 610-640. Modified worms were generated by SunyBiotech.

### Microscopy of *C. elegans* embryos

Embryos from adult worms were dissected on a 22 × 50 mm coverslip (Coring Catalog # 2975-225) in 10µL of egg salts buffer (ESB) containing 15-µm polystyrene beads (Sigma) using two 22-gauge needles. For microtubule depolymerization assays, ESB was mixed with nocodazole to make a 20µM nocodazole solution or 1% DMSO. Samples were then mounted onto plain 25 x 75 x 1 mm microscope slides (Fisher # 12-544-4). Time-lapse images were acquired with an inverted Nikon Eclipse Ti2-E microscope with a Yokogawa confocal scanner unit (CSU-W1), piezo Z stage, and an iXon Ultra 888 EMCCD camera (Andor), controlled by Nikon Elements software. For one cell embryos and individual late-development embryos, we used a 60X 1.2 NA Plan Apochromat water-immersion objective to acquire 41 × 0.5-µm Z-stacks with 100 ms exposures every min for 10 min starting slightly prior or during pronuclear meeting. 488 nm excitation (15 % laser power) was recorded using 2 × 2 binning followed by DIC imaging (92.3% iris intensity) using 1 x 1 binning.

### *In vitro* condensate assays

To assess self-assembly, GFP-labeled SPD-5 variants were incubated in condensate assembly buffer (25 mM HEPES, 150mM KCl, 0.5 mM DTT, 7.5% (w/v) PEG-3350). To assess recruitment, 1000 nM SPD-5::RFP was incubated in condensate assembly buffer for 2 min, then 10 nM SPD-5::GFP (final concentration) was added. All condensates were transferred to a pre-cleaned glass bottom imaging plate (Corning ref#4580; 96 well) and settled for 5 min before imaging with a Nikon Eclipse Ti-2E spinning disk confocal microscope (described in the live-cell imaging section) and either a 40X 1.25 NA silicone or 100X 1.35 NA silicone immersion objective.

### Circular dichroism

Pure SPD-5(a.a. 541-677) was dialyzed against Working Buffer (5 mM sodium phosphate pH 7.4, 150 mM NaF). Protein was diluted to 0.21 mg/mL, placed in a quartz cuvette with 0.1 cm path-length, then analyzed using a Jasco J-815 CD spectrometer. Data were accumulated 10 times, averaged, then analyzed with CONTIN as implemented on the web server DichroWeb using reference set 4.

### Embryo immunofluorescence

Prior to immunostaining, embryos were plated on 0.1mM IPTG plates with seeded with *perm-1(RNAi)* bacteria for 24 hr at 25°C to permeabilize the eggshells and inactivate the *or213ts* allele. Embryos were mounted in ESB (118mM NaCl, 48mM KCl, 2mM CaCl_2_, 2mM MgCl_2_, 25mM HEPES, balanced to 340 mOsm using sucrose) over Superfrost Plus slides (Fisher) and frozen in liquid nitrogen, fixed with methanol at −20°C for 10 min, then washed twice with TBST for 5 min. Samples were blocked in TBST +3% BSA for 30 min at room temperature, then incubated 1 hr with blocking buffer + primary antibody (1:5000 rabbit anti-SPD-5(lot 785))(Hamill et al., 2002). Slides were washed 3 times for 5 min with TBST, incubated for 1 hr in blocking buffer + secondary antibody (1:1000 donkey anti-rabbit alexa555 (Invitrogen), then washed 3 times for 5 min with TBST. Cells were mounted in Vectashield mounting medium with DAPI (Vector laboratories) and imaged by fluorescence confocal microscopy.

### Image quantification and statistical analyses

Images were analyzed using semi-automated, threshold-based particle analysis in FIJI. Data were plotted and statistical tests were performed using GraphPad prism. The sample size, measurement type, error type, and statistical test are described in the Figure legends where appropriate.

## SUPPLEMENTAL DATA

**Supplemental data set 1. Cumulative list of interacting pairs identified by XL-MS.**

See attached .xls file.

**Supplemental data set 2. Phospho-sites detected in purified SPD-5 by MS.** See attached .xls file.

**Supplemental data set 3. XL-MS peptide analysis of SPD-5(a.a. 541-677).** See attached .xls file.

**Supplemental data set 4. XL-MS peptide analysis of SPD-5(full-length).** See attached.xls file.

## Supplemental Tables

**TABLE S1.** Constructs for protein expression.

| protein | sequence | plasmid name | parent plasmid | N-term tag | C-term tag |
| --- | --- | --- | --- | --- | --- |
| SPD-5 | Full-length | JWV2 | pOCC28 | MBP-PreScission | PreScission-6xHis |
| SPD-5::GFP | Full-length | JWV1 | pOCC27 | MBP-PreScission | **eGFP**-PreScission-6xHis |
| SPD-5::RFP | Full-length | JWV3 | pOCC25 | MBP-PreScission | **tagRFP**-PreScission-6xHis |
| SPD-5::GFP | (R592K) | JWV13 | pOCC27 | MBP-PreScission | **eGFP**-PreScission-6xHis |
| SPD-5 | a.a. 541-677 | JWB127 | 6xHis_1B | 6xHis-TEV |  |
| SPD-5 | a.a. 541-677 (R592K) | JWB153 | 6xHis_1B | 6xHis-TEV |  |
| PLK-1(CA) | T194D | JWV11 | pOCC7 | 6xHis-PreScission |  |
| PLK-1(KD) | K67M | JWV12 | pOCC7 | 6xHis-PreScission |  |

**TABLE S2.**
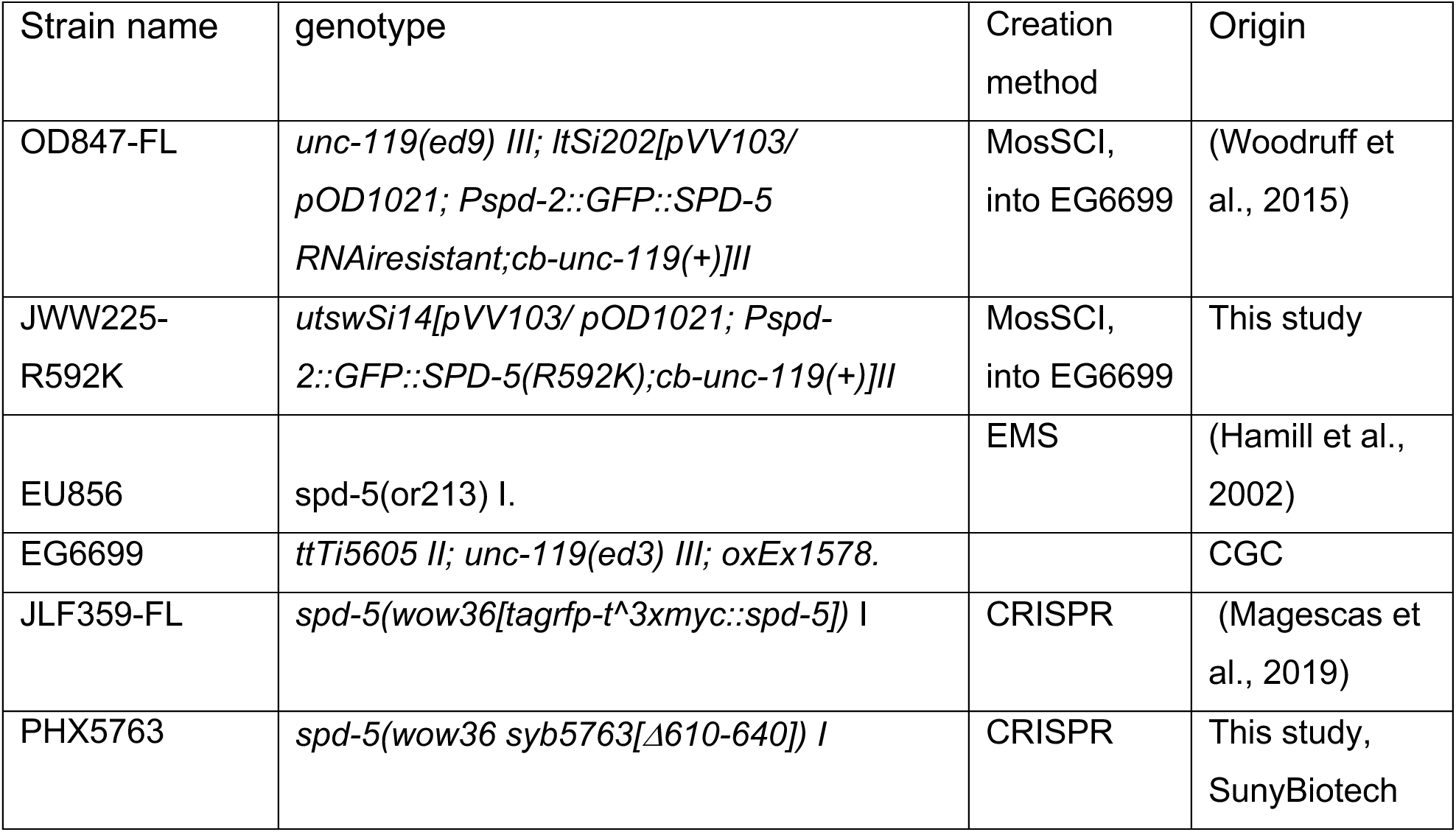
*C. elegans* strains.

## SUPPLEMENTAL FIGURE LEGENDS

**Figure S1.**
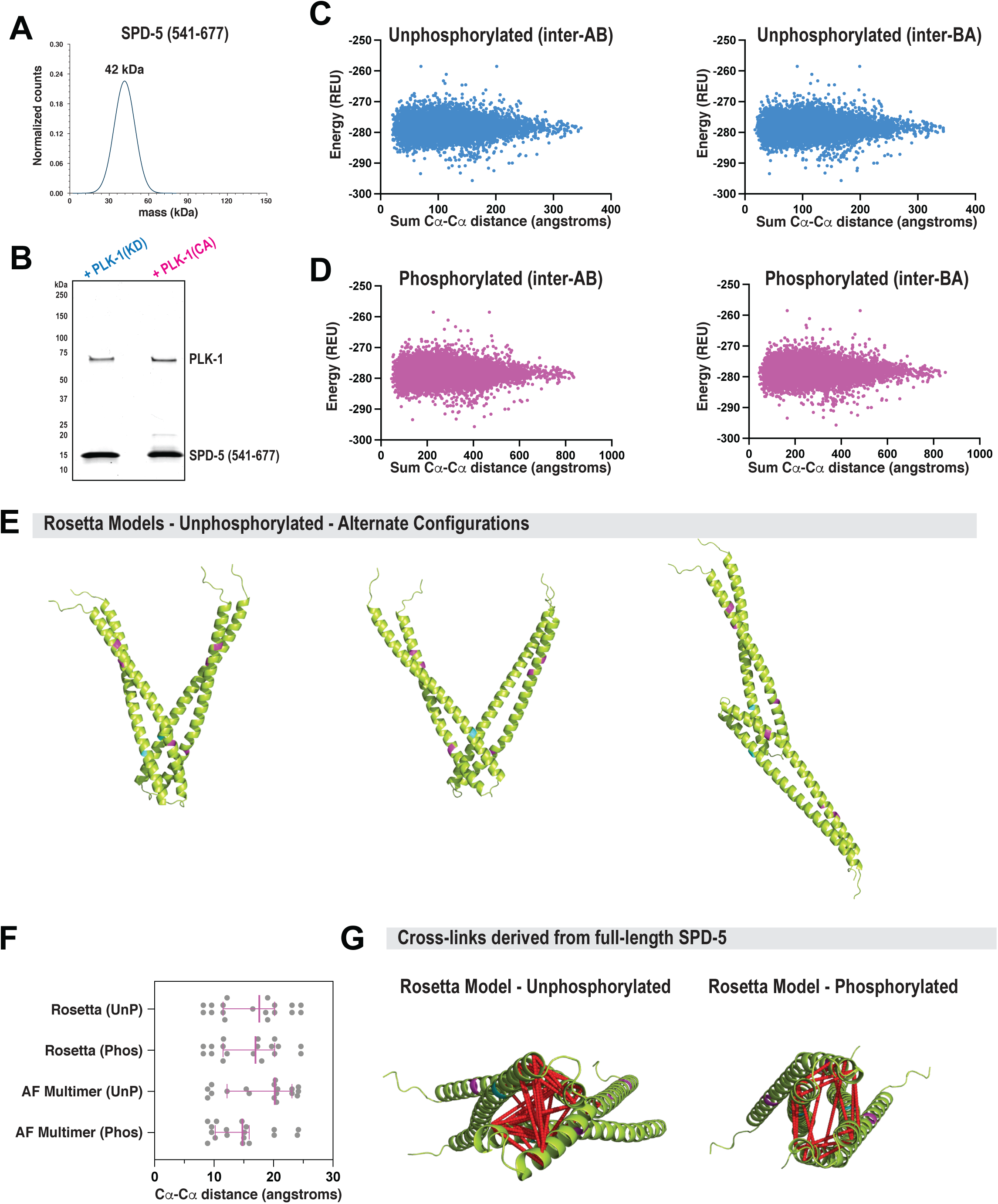
Biochemical and structural analysis of SPD-5(541-677). A. Mass photometry of 50 nM SPD-5(541-677). B. SPD-5(541-677) was incubated with PLK-1(KD) or PLK-1(CA) and analyzed by SDS-PAGE. The slower migrating bands (∼20 kDa) represent phosphorylated species of SPD-5. C. Evaluation of 10,000 models built in Rosetta for unphosphorylated SPD-5(541-677). D. Evaluation of 10,000 models built in Rosetta for phosphorylated SPD-5(541-677). E. Comparison of top-ranking alternative models for unphosphorylated SPD-5(541-677). F. Distances between alpha carbons of cross-linked residues in Rosetta and AlphaFold multimer models. Mean +/- 95% C.I. (N = 18 cross-links per sample). G. Evaluation of Rosetta structural models using cross-links (red dashed lines) derived from full-length SPD-5 in the presence of PLK-1(CA) or PLK-1(KD)(related to Figure 1B). Only cross-links that appeared in 2 or more samples are visualized.

**Figure S2.**
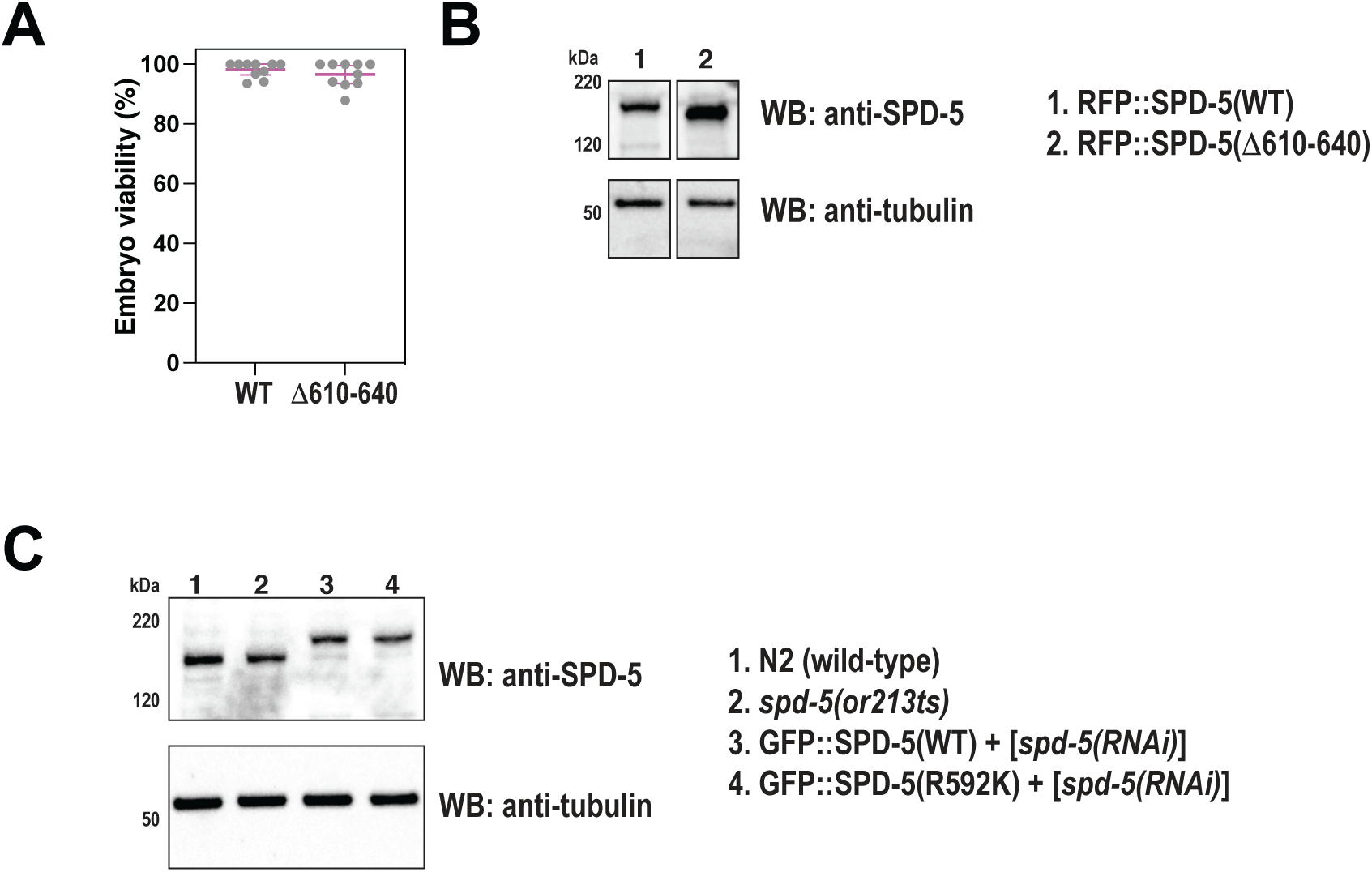
Embryo viability and protein expression controls. A. Viability of offspring in worms expressing wild-type (WT) or mutant (Δ610-640) RFP::SPD-5. Mean +/- 95% C.I. (N = 10 worms). Related to Figure 3. B. Western blots depicting expression levels of indicated proteins. Alpha tubulin was detected as a loading control. Related to Figure 3. C. Western blots depicting expression levels of indicated proteins. Alpha tubulin was detected as a loading control. Related to Figure 4.

**Figure S3.**
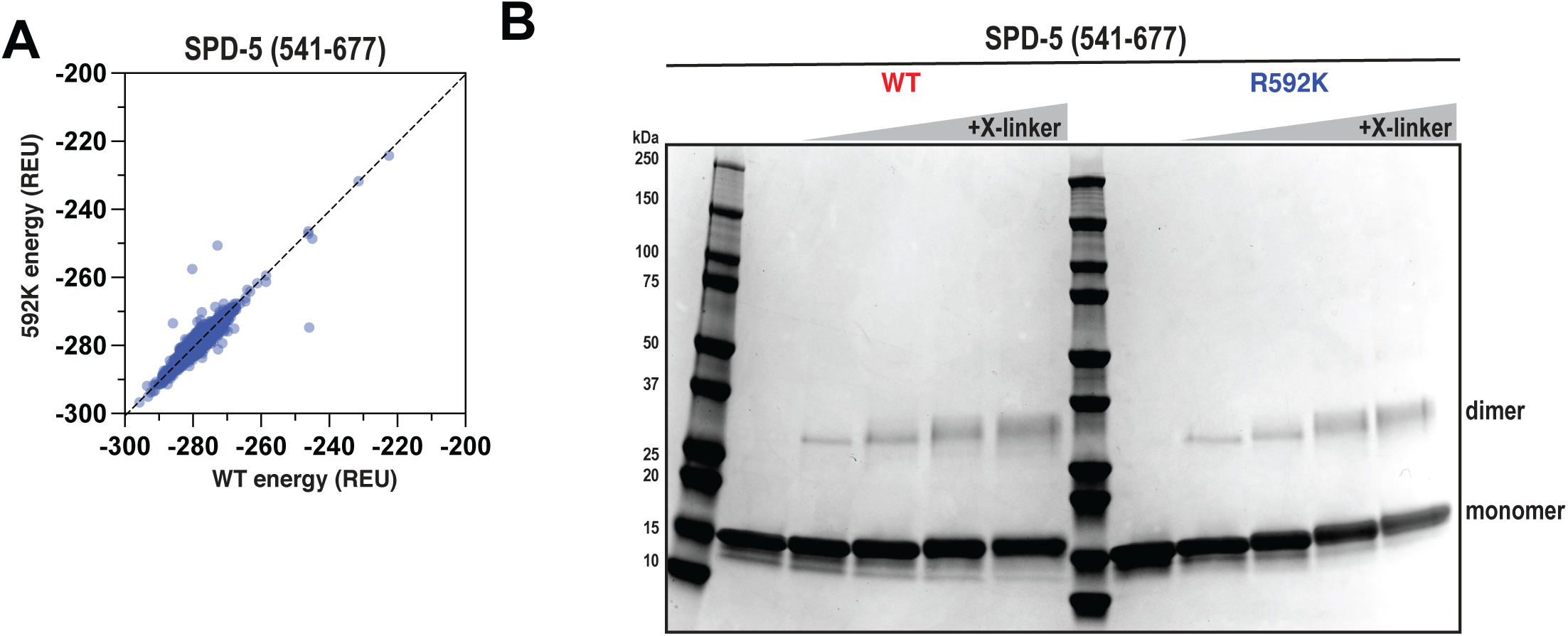
Biochemical analysis of the R592K mutation in SPD-5. A. The energetics of 10,000 *ab initio* models were calculated, then compared with the same model containing the R592K substitution. Each dot represents one model. Data on the diagonal indicate that the mutation does not change the energetics of folding in a particular model. B. SPD-5(541-677) (WT or R592K) was incubated with different amounts of DMTMM crosslinker (0-10 mM) for 45 min, then analyzed by SDS-PAGE.

## REFERENCES

Cabral, G., T. Laos, J. Dumont, and A. Dammermann. 2019. Differential Requirements for Centrioles in Mitotic Centrosome Growth and Maintenance. Dev Cell. 50:355–366 e356.

Carvalho, A., S.K. Olson, E. Gutierrez, K. Zhang, L.B. Noble, E. Zanin, A. Desai, A. Groisman, and K. Oegema. 2011. Acute drug treatment in the early C. elegans embryo. PLoS One. 6:e24656.

Chaudhury, S., M. Berrondo, B.D. Weitzner, P. Muthu, H. Bergman, and J.J. Gray. 2011. Benchmarking and analysis of protein docking performance in Rosetta v3.2. PLoS One. 6:e22477.

Conduit, P.T., K. Brunk, J. Dobbelaere, C.I. Dix, E.P. Lucas, and J.W. Raff. 2010. Centrioles regulate centrosome size by controlling the rate of Cnn incorporation into the PCM. Curr Biol. 20:2178–2186.

Conduit, P.T., J.H. Richens, A. Wainman, J. Holder, C.C. Vicente, M.B. Pratt, C.I. Dix, Z.A. Novak, I.M. Dobbie, L. Schermelleh, and J.W. Raff. 2014. A molecular mechanism of mitotic centrosome assembly in Drosophila. Elife. 3:e03399.

Conduit, P.T., A. Wainman, and J.W. Raff. 2015. Centrosome function and assembly in animal cells. Nat Rev Mol Cell Biol. 16:611–624.

Decker, M., S. Jaensch, A. Pozniakovsky, A. Zinke, K.F. O’Connell, W. Zachariae, E. Myers, and A.A. Hyman. 2011. Limiting amounts of centrosome material set centrosome size in C. elegans embryos. Curr Biol. 21:1259–1267.

Doxsey, S.J., P. Stein, L. Evans, P.D. Calarco, and M. Kirschner. 1994. Pericentrin, a highly conserved centrosome protein involved in microtubule organization. Cell. 76:639–650.

Dumont, S., and T.J. Mitchison. 2009. Force and length in the mitotic spindle. Curr Biol. 19:R749–761.

Enos, S.J., M. Dressler, B.F. Gomes, A.A. Hyman, and J.B. Woodruff. 2018. Phosphatase PP2A and microtubule-mediated pulling forces disassemble centrosomes during mitotic exit. Biol Open. 7.

Erpf, A.C., L. Stenzel, N. Memar, M. Antoniolli, M. Osepashvili, R. Schnabel, B. Conradt, and T. Mikeladze-Dvali. 2019. PCMD-1 Organizes Centrosome Matrix Assembly in C. elegans. Curr Biol. 29:1324–1336 e1326.

Farhadifar, R., C.H. Yu, G. Fabig, H.Y. Wu, D.B. Stein, M. Rockman, T. Muller-Reichert, M.J. Shelley, and D.J. Needleman. 2020. Stoichiometric interactions explain spindle dynamics and scaling across 100 million years of nematode evolution. Elife. 9.

Feng, Z., A. Caballe, A. Wainman, S. Johnson, A.F.M. Haensele, M.A. Cottee, P.T. Conduit, S.M. Lea, and J.W. Raff. 2017. Structural Basis for Mitotic Centrosome Assembly in Flies. Cell. 169:1078–1089 e1013.

Frokjaer-Jensen, C., M.W. Davis, C.E. Hopkins, B.J. Newman, J.M. Thummel, S.P. Olesen, M. Grunnet, and E.M. Jorgensen. 2008. Single-copy insertion of transgenes in Caenorhabditis elegans. Nat Genet. 40:1375–1383.

Fu, J., and D.M. Glover. 2012. Structured illumination of the interface between centriole and peri-centriolar material. Open Biol. 2:120104.

Gray, J.J., S. Moughon, C. Wang, O. Schueler-Furman, B. Kuhlman, C.A. Rohl, and D. Baker. 2003. Protein-protein docking with simultaneous optimization of rigid-body displacement and side-chain conformations. J Mol Biol. 331:281–299.

Hamill, D.R., A.F. Severson, J.C. Carter, and B. Bowerman. 2002. Centrosome maturation and mitotic spindle assembly in C. elegans require SPD-5, a protein with multiple coiled-coil domains. Dev Cell. 3:673–684.

Hannak, E., M. Kirkham, A.A. Hyman, and K. Oegema. 2001. Aurora-A kinase is required for centrosome maturation in Caenorhabditis elegans. J Cell Biol. 155:1109–1116.

Jiang, X., D.B.T. Ho, K. Mahe, J. Mia, G. Sepulveda, M. Antkowiak, L. Jiang, S. Yamada, and L.E. Jao. 2021. Condensation of pericentrin proteins in human cells illuminates phase separation in centrosome assembly. J Cell Sci. 134.

Jumper, J., R. Evans, A. Pritzel, T. Green, M. Figurnov, O. Ronneberger, K. Tunyasuvunakool, R. Bates, A. Zidek, A. Potapenko, A. Bridgland, C. Meyer, S.A.A. Kohl, A.J. Ballard, A. Cowie, B. Romera-Paredes, S. Nikolov, R. Jain, J. Adler, T. Back, S. Petersen, D. Reiman, E. Clancy, M. Zielinski, M. Steinegger, M. Pacholska, T. Berghammer, S. Bodenstein, D. Silver, O. Vinyals, A.W. Senior, K. Kavukcuoglu, P. Kohli, and D. Hassabis. 2021. Highly accurate protein structure prediction with AlphaFold. Nature. 596:583–589.

Kuhn, M., A.A. Hyman, and A. Beyer. 2014. Coiled-coil proteins facilitated the functional expansion of the centrosome. PLoS Comput Biol. 10:e1003657.

Laan, L., N. Pavin, J. Husson, G. Romet-Lemonne, M. van Duijn, M.P. Lopez, R.D. Vale, F. Julicher, S.L. Reck-Peterson, and M. Dogterom. 2012. Cortical dynein controls microtubule dynamics to generate pulling forces that position microtubule asters. Cell. 148:502–514.

Lawo, S., M. Hasegan, G.D. Gupta, and L. Pelletier. 2012. Subdiffraction imaging of centrosomes reveals higher-order organizational features of pericentriolar material. Nat Cell Biol. 14:1148–1158.

Lee, K., and K. Rhee. 2011. PLK1 phosphorylation of pericentrin initiates centrosome maturation at the onset of mitosis. J Cell Biol. 195:1093–1101.

Leitner, A., L.A. Joachimiak, P. Unverdorben, T. Walzthoeni, J. Frydman, F. Forster, and R. Aebersold. 2014a. Chemical cross-linking/mass spectrometry targeting acidic residues in proteins and protein complexes. Proc Natl Acad Sci U S A. 111:9455–9460.

Leitner, A., T. Walzthoeni, and R. Aebersold. 2014b. Lysine-specific chemical cross-linking of protein complexes and identification of cross-linking sites using LC-MS/MS and the xQuest/xProphet software pipeline. Nat Protoc. 9:120–137.

Lemaitre, R.P., A. Bogdanova, B. Borgonovo, J.B. Woodruff, and D.N. Drechsel. 2019. FlexiBAC: a versatile, open-source baculovirus vector system for protein expression, secretion, and proteolytic processing. BMC Biotechnol. 19:20.

Li, P., S. Banjade, H.C. Cheng, S. Kim, B. Chen, L. Guo, M. Llaguno, J.V. Hollingsworth, D.S. King, S.F. Banani, P.S. Russo, Q.X. Jiang, B.T. Nixon, and M.K. Rosen. 2012. Phase transitions in the assembly of multivalent signalling proteins. Nature. 483:336–340.

Magescas, J., J.C. Zonka, and J.L. Feldman. 2019. A two-step mechanism for the inactivation of microtubule organizing center function at the centrosome. Elife. 8.

Megraw, T.L., S. Kilaru, F.R. Turner, and T.C. Kaufman. 2002. The centrosome is a dynamic structure that ejects PCM flares. J Cell Sci. 115:4707–4718.

Mennella, V., B. Keszthelyi, K.L. McDonald, B. Chhun, F. Kan, G.C. Rogers, B. Huang, and D.A. Agard. 2012. Subdiffraction-resolution fluorescence microscopy reveals a domain of the centrosome critical for pericentriolar material organization. Nat Cell Biol. 14:1159–1168.

Mittasch, M., V.M. Tran, M.U. Rios, A.W. Fritsch, S.J. Enos, B. Ferreira Gomes, A. Bond, M. Kreysing, and J.B. Woodruff. 2020. Regulated changes in material properties underlie centrosome disassembly during mitotic exit. J Cell Biol. 219.

Moritz, M., M.B. Braunfeld, J.W. Sedat, B. Alberts, and D.A. Agard. 1995. Microtubule nucleation by gamma-tubulin-containing rings in the centrosome. Nature. 378:638–640.

Nakajo, M., H. Kano, K. Tsuyama, N. Haruta, and A. Sugimoto. 2022. Centrosome maturation requires phosphorylation-mediated sequential domain interactions of SPD-5. J Cell Sci. 135.

Ohta, M., Z. Zhao, D. Wu, S. Wang, J.L. Harrison, J.S. Gomez-Cavazos, A. Desai, and K.F. Oegema. 2021. Polo-like kinase 1 independently controls microtubule-nucleating capacity and size of the centrosome. J Cell Biol. 220.

Paix, A., A. Folkmann, D. Rasoloson, and G. Seydoux. 2015. High Efficiency, Homology-Directed Genome Editing in Caenorhabditis elegans Using CRISPR-Cas9 Ribonucleoprotein Complexes. Genetics. 201:47–54.

Pecreaux, J., J.C. Roper, K. Kruse, F. Julicher, A.A. Hyman, S.W. Grill, and J. Howard. 2006. Spindle oscillations during asymmetric cell division require a threshold number of active cortical force generators. Curr Biol. 16:2111–2122.

Pelletier, L., N. Ozlu, E. Hannak, C. Cowan, B. Habermann, M. Ruer, T. Muller-Reichert, and A.A. Hyman. 2004. The Caenorhabditis elegans centrosomal protein SPD-2 is required for both pericentriolar material recruitment and centriole duplication. Curr Biol. 14:863–873.

Rathbun, L.I., A.A. Aljiboury, X. Bai, N.A. Hall, J. Manikas, J.D. Amack, J.N. Bembenek, and H. Hehnly. 2020. PLK1- and PLK4-Mediated Asymmetric Mitotic Centrosome Size and Positioning in the Early Zebrafish Embryo. Curr Biol. 30:4519–4527 e4513.

Roostalu, J., N.I. Cade, and T. Surrey. 2015. Complementary activities of TPX2 and chTOG constitute an efficient importin-regulated microtubule nucleation module. Nat Cell Biol. 17:1422–1434.

Salisbury, J.L. 2003. Centrosomes: coiled-coils organize the cell center. Curr Biol. 13:R88–90.

Varadi, M., S. Anyango, M. Deshpande, S. Nair, C. Natassia, G. Yordanova, D. Yuan, O. Stroe, G. Wood, A. Laydon, A. Zidek, T. Green, K. Tunyasuvunakool, S. Petersen, J. Jumper, E. Clancy, R. Green, A. Vora, M. Lutfi, M. Figurnov, A. Cowie, N. Hobbs, P. Kohli, G. Kleywegt, E. Birney, D. Hassabis, and S. Velankar. 2022. AlphaFold Protein Structure Database: massively expanding the structural coverage of protein-sequence space with high-accuracy models. Nucleic Acids Res. 50:D439–D444.

Vasquez-Limeta, A., and J. Loncarek. 2021. Human centrosome organization and function in interphase and mitosis. Semin Cell Dev Biol.

Wang, Z., T. Wu, L. Shi, L. Zhang, W. Zheng, J.Y. Qu, R. Niu, and R.Z. Qi. 2010. Conserved motif of CDK5RAP2 mediates its localization to centrosomes and the Golgi complex. J Biol Chem. 285:22658–22665.

Wieczorek, M., S. Bechstedt, S. Chaaban, and G.J. Brouhard. 2015. Microtubule-associated proteins control the kinetics of microtubule nucleation. Nat Cell Biol. 17:907–916.

Woodruff, J.B. 2021. The material state of centrosomes: lattice, liquid, or gel? Curr Opin Struct Biol. 66:139–147.

Woodruff, J.B., B. Ferreira Gomes, P.O. Widlund, J. Mahamid, A. Honigmann, and A.A. Hyman. 2017. The Centrosome Is a Selective Condensate that Nucleates Microtubules by Concentrating Tubulin. Cell. 169:1066–1077 e1010.

Woodruff, J.B., and A.A. Hyman. 2015. Method: In vitro analysis of pericentriolar material assembly. Methods Cell Biol. 129:369–382.

Woodruff, J.B., O. Wueseke, and A.A. Hyman. 2014. Pericentriolar material structure and dynamics. Philos Trans R Soc Lond B Biol Sci. 369.

Woodruff, J.B., O. Wueseke, V. Viscardi, J. Mahamid, S.D. Ochoa, J. Bunkenborg, P.O. Widlund, A. Pozniakovsky, E. Zanin, S. Bahmanyar, A. Zinke, S.H. Hong, M. Decker, W. Baumeister, J.S. Andersen, K. Oegema, and A.A. Hyman. 2015. Centrosomes. Regulated assembly of a supramolecular centrosome scaffold in vitro. Science. 348:808–812.

Zhu, F., S. Lawo, A. Bird, D. Pinchev, A. Ralph, C. Richter, T. Muller-Reichert, R. Kittler, A.A. Hyman, and L. Pelletier. 2008. The mammalian SPD-2 ortholog Cep192 regulates centrosome biogenesis. Curr Biol. 18:136–141.

